# Phyloepigenetics in phylogeny analyses

**DOI:** 10.1101/2024.08.14.607911

**Authors:** Simeon Santourlidis

## Abstract

Long-standing, continuous blurring and controversies in the field of phylogenetic interspecies relations, associated with insufficient explanations for dynamics and variability of speeds of evolution in mammals, hint to a crucial missing link. It has been suggested that transgenerational epigenetic inheritance and the concealed mechanisms behind play a distinct role in mammalian evolution. Here, a comprehensive sequence alignment approach in hominid species, i.e., *Homo sapiens, Homo neanderthalensis, denisovan human, Pan troglodytes, Pan paniscus, Gorilla gorilla and Pongo pygmaeus*, comprising conserved CpG islands of housekeeping genes, uncover evidence for a distinct variability of CpG dinucleotides. Applying solely these evolutionary consistent and inconsistent CpG sites in a classic phylogenetic analysis, calibrated by the divergence time point of the common chimpanzee (Pan troglodytes) and the bonobo or pygmy chimpanzee (Pan paniscus), a "phylo-epigenetic” tree has been generated which precisely recapitulates branch points and branch lengths, i.e., divergence events and relations, as they have been broadly suggested in the current literature, based on comprehensive molecular phylogenomics and fossil records. I suggest here that CpG dinucleotides changes at CpG islands are of superior importance for evolutionary development and determine the emerging DNA methylation profiles.

## 1. Introduction

C.H. Waddington presented conceptual explanations for how the approx. 200 different cell types, despite all having the same genetic constitution, contribute each differently, within the shortest time, to the successful development of the evidently complex human fetus, individually adapted and prepared for the upcoming environmental conditions and challenges [1]. Notably, over the last 40 years, we gained comprehensive knowledge on the epigenetic mechanisms behind, in particular on DNA methylation, annotating and reorganizing the genome, involved in achieving directed, meaningful cell type specific functionality and phenotype. Hence, epigenetics has been proceeded to become of the similar fundamental importance as classical genetics and any branch of biology, e.g., evolution, and of medicine, e.g., cancer research, can only be better understood if epigenetics is appropriately taken into account.

However, it is an unsolved mystery on the nature of the epigenetic master initiator within the fertilized oocyte, inherited by the parental generation. He leads to the imposition of the specific epigenetic profile, evidently, evolutionary shaped by the experiences of many ancestral generations in intensive, daily confrontation with the challenging environmental conditions and requirements.

Nevertheless, currently the neo-Darwinian view of evolution with selection as its central pillar is still dominating over the upcoming conceptions on transgenerational inheritance of acquired traits which are closer to Lamarckian inheritance. The latter are controversially discussed and in particular when they are linked to postulated transgenerational inheritance of acquired DNA methylation patterns, since primordial germ cells essentially erase DNA methylation [2].

Regardless, the complete organism, originated in one zygote, ultimately testified versatile evolutionary developments and adaptations and this in different models, i.e., the different species. Here, what classical genetics mostly denied, the plasticity of epigenetics and its connection to the environment provide new explanations for the requested dynamic variations in evolutionary speeds to overcome and adapt to fluctuant and challenging environmental conditions.

Currently, the ‘molecular-clock principle’ is widely used to assess the timescale of organismal evolution [3], to infer time and correlation between lineage divergence time and concurrent environmental changes [4]. Its assumption is that the number of molecular character changes strikingly reflects phylogenetic distance of organisms and therefore allows the inference of the exact time point since the divergence of species. In addition, simplified, equal evolutionary rates are assumed between the species to be compared.

On the other hand the scientific literature reveals yet unexplained, impressive discrepancies since molecular time-scales frequently have suggested that lineage origination may be twice as old as fossils imply. For example metazoan phyla have been originated several hundred million years (ca. 750–1200 Ma) before the Cambrian explosion (ca. 560 Ma) of these animals in the fossil record. And further, most orders of birds and mammals appear to have originated within the mid-late Cretaceous (ca. 80–100 Ma), while the fossil record proposes a sudden appearance of modern lineages in the early Tertiary (65–54 Ma) [4]. Interestingly in this context the authors conclude that effects of the environment on shaping bird and mammal biodiversity through time are optimally testable for diversification, not origination and pointed out the time of diversification within avian order and family appears to correlate with the two largest environmental changes in Tertiary history [4].

Thus, evolutionary tracking based solely on genetic variation remains difficult and blurred. Understanding the manifestation and role of epigenetic inheritance could lead to new approaches to more accurately track evolutionary processes.

The most studied epigenetic mechanism is DNA methylation, which occurs at CpG dinucleotides and is involved in cell-type specific epigenetic suburbanization of the genome in active, silent and transcriptional competent parts. It participates in the control of the outcome of the genetic content of a particular cell. More than 60% of human genes possess a 0.4-2 kb long, CpG rich 5’-gene area of regulatory importance, surrounding the transcriptional start site, which is called a ‘CpG-island’. In particular, these CpG-islands at housekeeping genes remain unmethylated throughout development and in all tissues, which is thought to protect them from erosion [5]. Full and partial methylation at specific, single CpG dinucleotides within these CpG islands affect gene expression [6]. In contrast to the spontaneous occurring hydrolytic cytosine deamination which leads to uridine in DNA and is removed by uracil-DNA glycosylase, the same reaction at methylcytosin yields the normal DNA base thymine which largely persists at the affected positions and this results in increased mutation rates of CpG to TpG and CpA, respectively, in both strands, due to the palidromy of the CpG dinucleotide and the methylation maintenance property of the DNA-methyltransferase 1 (DNMT-1) [7]. The transition rate of methylated CpG to TpG is 10–50 times higher than other transitional changes [8]. From this it would generally be assumed that steadily unmethylated CpGs within CpG islands of conserved genes should also remain evolutionary conserved.

In contrast, it has been suggested that CpGs seem to be hotspots of mutations related to speciation [9]. The authors postulated that, on the one hand, certain CpG-related mutations would promote genomic

‘flexibility’ in evolution, i.e., the ability of the genome to expand its functional possibilities; on the other hand, CpG-related mutations in SNPs would relate to genomic ‘specificity’ in evolution, thus, representing mutations that would associate with phenotypic traits relevant for speciation [9]. Furthermore, within orthologous CpG islands, those distinctive regulatory regions, associated with up to two thirds of vertebrate gene promoters, CpG dinucleotides differ greatly in terms of base composition and frequency among vertebrate species, contributing to a faster evolution than this of CpG-poor promoters [6].

Going in a similar direction I hypothesized that alterations on CpG dinucleotides would play a distinct evolutionary role, by direct comparison to other nucleotide changes, firstly due to their clearly increased mutation rate when methylated, and secondly by their broadly proven functional relevance for gene expression. If so, evolutionary changes within certain CpGs, considered separately from all other nucleotide positions which undergo changes in lower rates, may reflect interspecies evolutionary relationships in a more enhanced resolution as conventional molecular analyses do, and presumably, this could be preferentially become visible if closer related species would be analyzed [10]. Further I assumed, in analogy to interspecies single nucleotide variations that CpG dinucleotides should vary lesser between closely related species, which have been diverged recently, as opposed to cases of earlier divergence. Thus, are CpG dinucleotides of a distinct role for evolution, and may they therefore harbor and provide an improved phyloepigenetic tool to more accurately assess phylogeny relations? When yes, how to prove it and how to explain their role within a requested epigenetic transgenerational inheritance, since at least in generational scale erasure of methylation in the germ line has been proven?

To test this hypothesis and especially to find a suitable paradigm which might demonstrate the distinct importance of CpG mutations I developed the following experimental approach. I chose two dozen conserved housekeeping genes, picked up their CpG islands and I lined them up one after the other. I aligned them between a selection of close related species from which I can assume that they evolve at a comparable speed. I compare and evaluate all conventional non CpG mutations versus the CpG transitions and transversions. I assumed that this highly conserved scenario might reveal even slight differences in the relative frequency of the non CpG versus the CpG mutations and furthermore their different significance for the manifestation of the species relationships. If such differences exist they should penetrate here and become visible in appropriate phylogenetic tree analyses. The chosen species were primates, i.e. homo, gorilla, chimpanzee, bonobo, orangutan and to get hopefully a more reliable calibration point, in addition, the altai neanderthal and denisova genomes were chosen [11].

## 2. Materials and Methods

Conserved housekeeping genes were randomly chosen from a list provided by Tel Aviv University server at https://www.tau.ac.il/~elieis/HKG/ and based on the publication of Eisenberg et al. on human housekeeping genes [12]. The 5′regions of the following two dozen genes of Homo sapiens were picked up from the nucleotide database of the national library of medicine at https://www.ncbi.nlm.nih.gov/. The CpG islands of these genes were lined up one behind the other in this order: ACTR1A, CALR, AKIRIN1, H3-3B, BTF3, EED, AKIRIN2, AP1B1, HUS1, ITCH, SIRT2, PUM1, HAT1, HDAC2, H1FX, TUBB, PCNA, PNN, POLE3, POMP, SMU1, TOX4, NCL, RING1. The homologous primate sequences of chimpanzee, bonobo, gorilla, and orang-utan derived from BLAT search from the assemblies, panTro6 (chimp), panPan3 (bonobo), gorGor6 (gorilla), ponAbe3 (orang-utan) at https://genome.ucsc.edu/cgi-bin/hgBlat (accessed on 10 October 2023) [13]. The homologous altai neanderthal and denisova sequences derived from genome browser, JBrowse 1.12.1, at https://bioinf.eva.mpg.de/jbrowse/ (accessed on 4 November 2023) [11]. All CpG rich 5′regions were inspected by Repeatmasker at https://www.repeatmasker.org/, to exclude repetitive DNA elements, e.g., LINE-1, Alu etc.

I aligned all these sequences containing the lined-up CpG islands one behind the other by the multiple alignment program for nucleotide sequences MAFFT (v7.511) at https://mafft.cbrc.jp/alignment/server/ using the default settings, except scoring matrix for nucleotide sequences: 1PAM / κ=2, when aligning closely related DNA sequences and I use the iterative refinement method FFT-NS-i [14, 15]. For reconstructing of the phylogenetic tree, I used UPGMA (unweighted pair group method with arithmetic mean) at https://mafft.cbrc.jp/alignment/server/phylogeny.html, which is considered to be a simple agglomerative hierarchical clustering method which assumes a constant rate of evolution (molecular clock hypothesis) [16]. The analyses were done after replacement of all CpGs, which are consistent and part-consistent in all CpG rich sequences in all seven species. Here every CpG, consistent in all orthologous sequences was replaced by an “A” and every same positioned dinucleotide in every species of a position with at least one CpG in any of the other species was replaced by a “T” (Fig. 1 B). Visualization and comparing of the phylogenetic trees was done by phylo.io at http://phylo.io/ [17]. Percent Identity Matrix was created by Clustal 2.1 at https://www.ebi.ac.uk/ [18].

**Figure 1.**
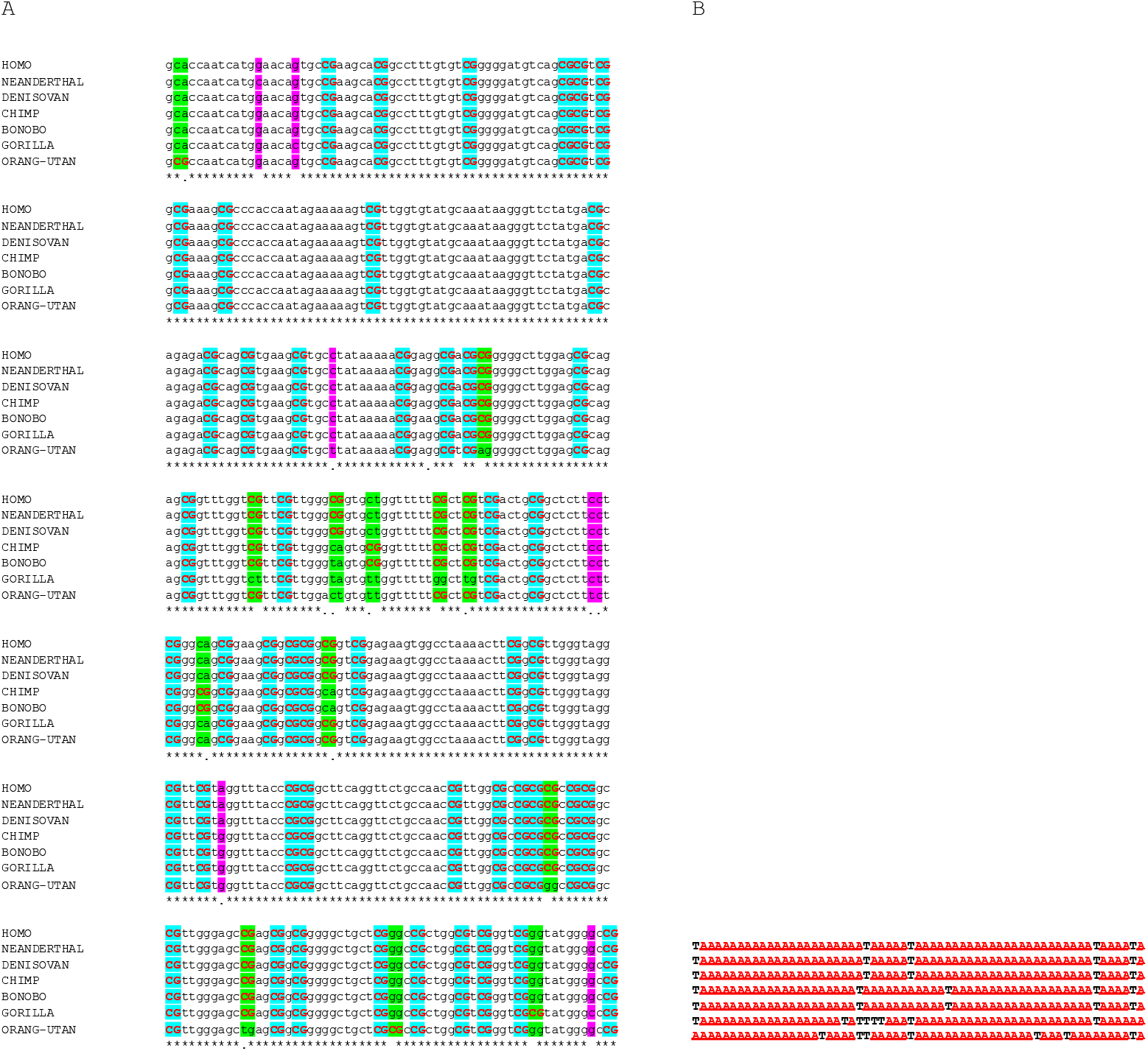
**A)** Representative excerpt of the alignment of assembled 24 5′ CpG rich sequences (CpG islands) of conserved housekeeping genes in hominids. All single nucleotide polymorphisms (SNPs) outside CpGs present in at least one of the seven species are highlighted in purple (approx. 4.8% of all possible, lined up dinucleotide positions). All CpG dinucleotides largely preserved but affected by at least one nucleotide alteration in at least one of the species are highlighted in green (that is approx. 3.2% of all possible, lined up dinucleotide positions and 17 % of all CpG positions). All CpG dinucleotides consistently preserved in all species are highlighted in light blue. CpGs are highlighted in red. **B)** Representative excerpt of the corresponding A/T aligment in which only the CpG positions are depicted, i.e., all CpGs, including those consistent in all seven species were replaced by “A” and those altered in least at one of the species were replaced by a “T”.

I chose the divergence time point of the common chimpanzee (Pan troglodytes) and the bonobo or pygmy chimpanzee (Pan paniscus), that is 1.7 My [22], as the calibration point for my phylogeny trees of Fig. 2.

**Figure 2.**
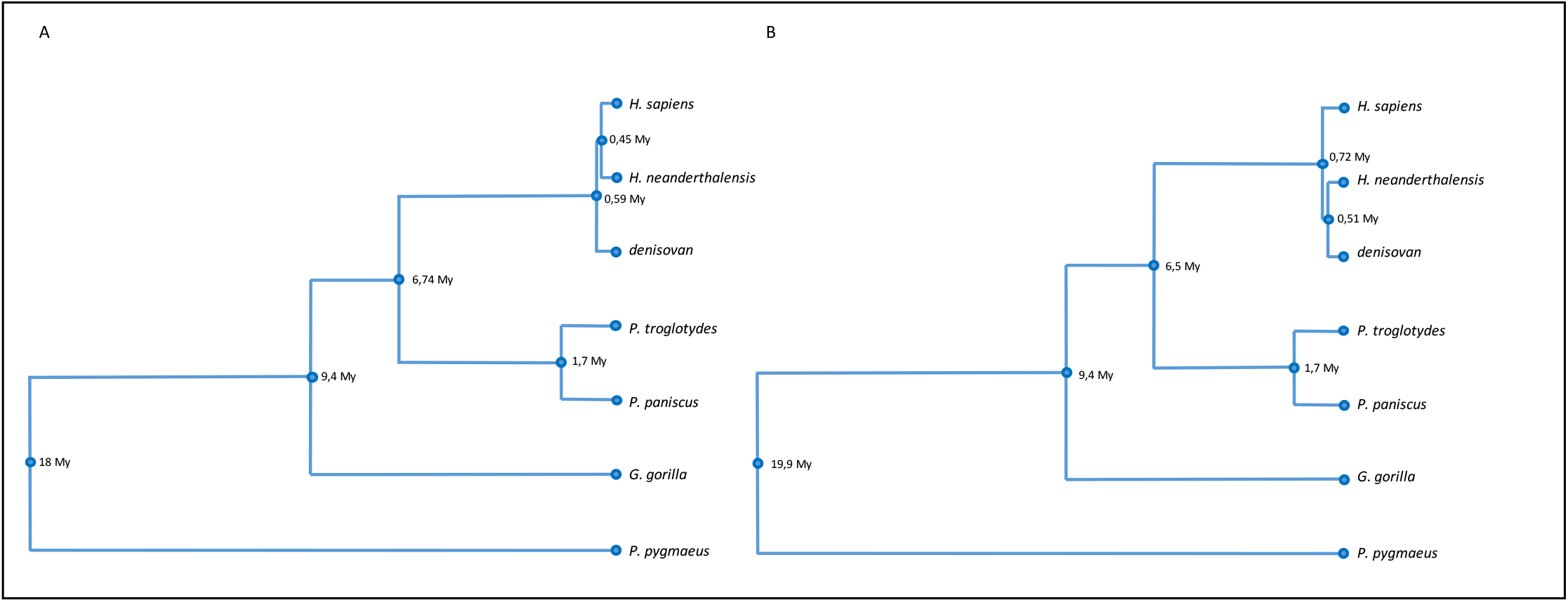
Phylogenetic (**A**) and “phylo-epigenetic” (**B**) relations between 7 primate species based on genetic and epigenetic, respectively, differences of CpG islands of 24 conserved housekeeping genes. The phylogram of the left panel displays the phylogenetic relation of these primate species, based on single nucleotide changes including CpG sites, small deletions and nucleotides gains. The phylogram of the right panel displays the “phylo-epigenetic” relation of these species, based on all consistent occurring CpG dinucleotides and all CpG dinucleotide differences of these CpG islands of conserved housekeeping genes.

## 3. Results

An alignment of the randomly chosen 24 CpG rich, lined up 5’-regulatory regions of conserved housekeeping genes from all seven species reveals the grade of conservation (an excerpt of this alignment of 420 nt is presented in Fig.1 A). The sequences aligned were of a total length of 27061 nucleotides from Homo sapiens, 27061 from Homo neanderthalensis, 27060 from denisovan human, 27037 from Pan troglodytes, 27072 from Pan paniscus, 27040 from Gorilla gorilla and 27037 from Pongo pygmaeus. The sequence homologies are for example 99.9% between Homo sapiens, Homo neanderthalensis and denisovan human. The differences between these species and Pan troglodytes, Pan paniscus, Gorilla gorilla and Pongo pygmaeus are 99 %, 99 %, 98.6 % and 97.3 %, respectively. All sequence homologies between these seven sequences are listed in detail in %, in table 1. The largest difference appear between the hominins and Pongo pygmaeus with 97.18 %.

**Table 1.** Percent Identity Matrix for 24 CpG rich 5’-regions of conserved housekeeping genes of all seven hominid species.

|  | Human | Neanderthal | Denisovan | Chimp | Bonobo | Gorilla | Orang-utan |
| --- | --- | --- | --- | --- | --- | --- | --- |
| Human | 100 | 99.90 | 99.89 | 99.01 | 99.01 | 98.63 | 97.32 |
| Neanderthal | 99.90 | 100 | 99.90 | 98.98 | 98.98 | 98.60 | 97.29 |
| Denisovan | 99.89 | 99.90 | 100 | 98.99 | 98.99 | 98.62 | 97.31 |
| Chimp | 99.01 | 98.98 | 98.99 | 100 | 99.74 | 98.56 | 97.23 |
| Bonobo | 99.01 | 98.98 | 98.99 | 99.74 | 100 | 98.57 | 97.23 |
| Gorilla | 98.63 | 98.60 | 98.62 | 98.56 | 98.57 | 100 | 97.18 |
| Orang-utan | 97.32 | 97.29 | 97.31 | 97.23 | 97.23 | 97.18 | 100 |

Thus, these sequences of lined up 24 CpG rich 5′-regions (CpG islands) of conserved housekeeping genes are highly conserved among these hominid species, with sequence homologies between 99.9 % and 97.2 %.

All single nucleotide positions outside CpGs, mutated at least in one of the primate sequences within the whole alignment are ca. 2.4 % of the ca. 27060 nucleotides. In order to compare the mutation rate of mutations at non CpG dinucleotides with those at CpG dinucleotides, the ca. 27060 nucleotides of the entire sequence were divided by 2 and all lined up dinucleotides were again inspected for mutations. Thus, 1.2% of those ca. 13530 dinucleotide positions were mutated at non CpG positions at least in one of the species. 3.1 % of all these, 13530 dinucleotide positions were mutated at orthologous dinucleotide positions where a CpG is present in at least one of the species. We have in total 2495 CpG dinucleotide positions within all possible lined up 13530 dinucleotide positions and of those 17 % were altered at least in one of the species. 2080 CpG dinucleotide positions were conserved throughout all species.

By counting, the following numbers of transitions and transversions were found, that they are present at dinucleotide positions which show one CpG dinucleotide in at least one of the species.: CG to TG 127 (5.1%), CG to CA 109 (4.4%), CG to GG 59 (2.4%), CG to CC 45 (1.8%), CG to CT 44 (1.8%), CG to AG 31 (1.2%). Regarding solely the CpG transitions to TpG and CpA from the perspective of every single species, table 2 depicts that the lowest number of 94 transitions at dinucleotide positions with a TpG or a CpA at that place in at least one of the other species are present in Gorilla gorilla and the highest number of 115 in Pongo pygmaeus. Of note, CpA corresponds to TpG in the complementary strand, which might result from a hydrolytic deamination of a methylated CpG. The corresponding numbers from all the other species are presented in table 2.

**Table 2.** The numbers of transitions to TpG and CpA in any of the species within the whole alignment from the perspective of every listed species with CpGs at the considered positions.

|  |  |
| --- | --- |
| homo | 98 |
| neanderthal | 101 |
| denisovan | 99 |
| chimp | 95 |
| bonobo | 95 |
| gorilla | 94 |
| orang-utan | 115 |

A representative excerpt of the alignment, by labelling illustrating the relevant, described positions and their changes is shown in Fig. 1A. For this part of the alignment, Fig 1B illustrates, the corresponding “A/T” alignment which refers exclusively to the throughout in all species conserved and non-conserved CpG positions (A) and to CpG dinucleotide linked transitions and transversions (T), orthologically, equally placed with one CpG dinucleotide in at least one of the species (Fig. 1B). Thus, within the sequence conserved CpG islands of conserved housekeeping genes a significant alteration rate persists at CpG dinucleotides and hereby predominantly CpG to TpG and CpA transitions are present.

Finally, the UPGMA phylogenetic trees, once based on all single nucleotide changes including CpG sites, small deletions and nucleotides gains (Fig. 2A) and one based sole on consistent CpG dinucleotides and CpG alterations as explained (Fig. 2B), preferably refer to here as phylo-epigenetic tree, are presented (Fig.2). The first one displays a closer relation of Homo sapiens to Homo neanderthalensis than to denisovan human. All the other branchpoints and branches indicate divergence events and species relations, respectively, as they are established in the literature. The tree of Fig. 2B resembles the phylogenetic tree of Fig. 2A, with the difference that Homo neanderthalensis appear nearer to denisova human than to Homo sapiens. When both trees are calibrated by the divergence time point of the common chimpanzee (Pan troglodytes) and the bonobo or pygmy chimpanzee (Pan paniscus), that is 1.7 My [22], the phylo-epigenetic tree displays evolutionary distances between the species as they have been suggested [19,20]. Thus, the resulting phylo-epigenetic tree based solely on all CpG dinucleotides and CpG dinucleotide differences reveals an image of these interspecies relations that precisely fits to the current established knowledge in this field of research.

## 4. Discussion

The most phylogenetically distant great apes from humans with the most ancestral karyotype among all hominids, providing an informative perspective on hominid evolution are the orang-utan species [19]. The speciation of orang-utan, is thought to have occurred no earlier than the Middle Miocene (12-16 Myr ago), as fossil apes before that differ substantially from what we might expect of an early great ape [20]. Based on comprehensive statistical analyses of molecular data and proper fossil calibration, the divergence date for gorillas and the human/chimpanzee clade has been suggested to range from 7-9 MYA and for humans and chimpanzees from 4-6 MYA. This is generally compatible with the known primate fossil record or recent molecular studies [21].

The common chimpanzee (Pan troglodytes) and the bonobo or pygmy chimpanzee (Pan paniscus) represent the closest living hominid species to humans and the most-recently diverged ape species (around 1.7 million years ago) [22].

It has been shown that Neanderthals contributed genetically to modern humans outside Africa 47,000-65,000 years ago, and it has been proposed that the ancestors of Neanderthals from the Altai Mountains and early modern humans met and interbred many thousands of years earlier than previously thought [23]. The analysis of a Neanderthal genome from a cave in the Altai Mountains in Siberia suggests they diverged 550,000-765,000 years ago. The analysis of a Denisovan genome from the same cave in the Altai Mountains further suggests that Neanderthals and Denisovans diverged 381,000–473,000 years ago [23]. The main result of this study is that for the evolution of these hominid species a "phylo-epigenetic tree” has been built up, which based solely on evolutionary consistent and inconsistent CpG sites (Fig. 2B). Noteworthy, it precisely recapitulates branch points and branch lengths, i.e., divergence events and rates of genetic changes, relations, as they have been broadly proposed and constitute the current state of knowledge in this research field, based on comprehensive molecular phylogenomics and fossil records [19,20]. This demonstrates the overriding importance of these CpG sites which are capable to bear the gene regulating, epigenetic mark, methylation, for transgenerational epigenetic inheritance.

Within this evolutionary comparison of hominids, even in highly conserved housekeeping genes with conserved sequences of CpG islands, the individual CpGs show a considerable dynamic rate of mutations which are predominantly CpG to TpG and CpA transitions. A possible explanation might be, that individual methylated CpG positions within these conserved CpG islands, had been preserved during the demethylation events of the early preimplantation developmental stage to become afterwards subjected to spontaneous hydrolytic deamination. This has occurred before the primordial germ cells of the new generation are separated. From then on these mutations persist within the lineage, engaging the positions of former CpG dinucleotides, which presumably, originally, decisively had impacted gene expression in dependence to their methylation state. Hence, these mutations should have an altering impact on the regulation of these genes. It has been already proposed by others that a stochastic, incomplete removal of DNA methylation marks takes place during a window of opportunity in the zygote and early embryo and that genes affected are sensitive to dosage and this may be associated with evolutionary advantages [8]. Others have suggested that mutations which globally affect epigenetic marking and expression variability are potentially advantageous in a variable environment [24]. That is, in respect of methylated CpG linked hydrolytic deamination events, opposed to the fact, that a complete deletion of DNA methylation in the primordial germ cells has been demonstrated [2]. I conclude here that this is due to methodological limitations to detect occasional events of persisting punctual methylation, stochastically and infrequently occurring in an evolutionary timescale, or/and hydrolytic deamination has already altered methylated CpGs, before separation of the primordial germ cells, thus masking them.

The work of hundreds of laboratories has established CpG islands as distinctive regulatory regions within the vast excess of genomic DNA sequence [6]. It is broadly accepted that punctual DNA methylation influences gene expression by the spatial impediment of transcription factor binding sites for their accessibility and by influencing DNA impact on nucleosomal positioning around the transcription start sites [6]. It is concluded here that substitution of a methylated cytosine by a thymine at specific, relevant CpG positions would also have a decisive impact on regulation of specific genes.

Given that we are looking at conserved housekeeping genes which prepossess an essential and fundamental role for the phenotype and function of every cell, we have to conclude that subtle evolutionary changes at these CpG dinucleotides might have a significant impact on the traits associated with these genes.

Further, I hypothesize that beyond these randomly chosen housekeeping genes, comparable analyses like this one, would reveal within the CpG islands of certain individual genes, distinct CpG constellations and profiles, respectively, which are acting as epigenetic hotspots of evolution (EHE). I expect significant enhanced, evolutionary CpG alteration rates of these EHE. This I presume due to their distinct, basic, functional relevance associated with phenotypic traits in connection with evolutionary fluctuating pressures of selection. In this respect, I am eager to see comparable studies like this on differentially methylated CpG islands which are essential for the dynamic embryonic development, e.g., OCT4, and tissue-specific expression profiles, e.g., MyoD. They constitute 5% of all CpG islands [6].

I suggest that a part of epigenetic inheritance is provided by effective turn over mechanisms, which in an evolutionary timescale are able to bypass early embryonic epigenetic resetting and lead to mutations at regulatory important CpG dinucleotides which former had differentially impacted gene regulation in dependency of their methylation state. In this scenario, it has to be requested that until now unknown mechanisms should be able to restore in evolutionary time scales new CpG dinucleotides at exact these relevant positions. Summing up, I postulate that evolutionary shaped fluctuation at exactly these important CpG sites is of superior importance for evolutionary plasticity. In this context it is accentuated that Bernhard Horsthemke and Adrian Bird refer recently [2] to the interesting study of Takahashi et al. [24] which shows that the privileged immunity to DNA methylation of a CpG island can be seriously compromised by transient local alteration of its DNA sequence and an acquired aberrant methylation pattern can be transmitted across generations. In their concluding remarks they pointed out that we now need to understand the molecular mechanisms that underlie this abrupt change of epigenetic status. Importantly, the relevance of this model system to acquisition and transmission of naturally occurring epigenetic variation has yet to be established [2].

It can be postulated that, on an evolutionary scale, the selective alteration of individual CpGs in the CpG islands decisively changes the profile of these distinctive regulatory regions with regard to their immunity to methylation. This would result in significant changes in the gene regulation of the genes involved. A provocative conclusion at that point would be that precisely these changing CpG positions at all genome wide regulatory CpG positions represent an epigenetic master initiator who determines the inherited and by the environment shaped genome-wide DNA methylation profile established in the early embryonic development. It is hidden in the DNA, what is to happen on it later.

Finally, similar phylo-epigenetic analyses, will be conducted, supported by an appropriate algorithm, comprising substantial larger parts of the genome, to uncover to what extent the approach presented here can further dissect phylo-epigenetic relationships in detail.

## 5. Conclusions

I conclude that, in evolutionary time scales, changes in differentially methylatable CpG dinucleotides within CpG islands play a superior role for evolutionary developments. It is postulated that these changes are successfully inherited through the germ line, determining emerging methylation profiles and are a central component of the transgenerational epigenetic inheritance.

## 6. Declarations

### Ethics approval and consent to participate

Not applicable

### Consent for publication

Granted

## Availability of data and materials

Data of this study are available from S.S. on request

## Authors’ contributions

S.S.; conceptualization, data generation, data analyses and interpretation, original draft preparation, figures preparation, writing—review and editing.

## Acknowledgements

I thank the Heinrich-Heine-University for my studies and my employment. I thank Dr. Wolfgang Schulz for critical comments on the article. Simeon Santourlidis wishes to dedicate this work to his 3 grandchilds, Simeon, Trias, Christos and his daughter Alexandra and his wife Tzanetina.

## Authors’ information (optional)

Simeon Santourlidis has been an employee of Universitätsklinikum Düsseldorf (UKD), Heinrich-Heine-Universität, Düsseldorf since April 15, 1997 and is professor of molecular medicine.

